# Measurement of Cervical Neuronal Activity during Stress Challenge Using Novel Flexible Adhesive Surface Electrodes

**DOI:** 10.1101/2022.07.10.499484

**Authors:** Yifeng Bu, Jonas F. Kurniawa, Jacob Prince, Andrew K. L. Nguyen, Brandon Ho, Nathan L. J. Sit, Timothy Pham, Vincent M. Wu, Boris Tjhia, Andrew J. Shin, Tsung-Chin Wu, Xin M. Tu, Ramesh Rao, Todd P. Coleman, Imanuel Lerman

## Abstract

This study introduces a flexible, adhesive-integrated electrode array that was developed to enable non-invasive monitoring of cervical nerve activity. The device uses silver-silver chloride as the electrode material of choice and combines it with a novel electrode array consisting of a customized biopotential data acquisition unit and integrated graphical user interface (GUI) for visualization of real-time monitoring. Preliminary testing demonstrated this novel electrode design can achieve a high signal to noise ratio during cervical neural recordings. To demonstrate the capability of the surface electrodes to detect changes in cervical neuronal activity, the cold-pressor test (CPT) and a timed respiratory challenge were employed as stressors to the autonomic nervous system. This sensor system recording, a new novel technique, was termed Cervical Electroneurography (CEN). By applying a custom spike sorting algorithm to the novel electrode measurements, neural activity was classified in two ways: 1) pre-to-post CPT, and 2) during a timed respiratory challenge. Unique to this work: 1) rostral to caudal channel position-specific (cephalad to caudal) firing patterns and 2) cross challenge biotype-specific change in average CEN firing, were observed with both CPT and the timed respiratory challenge. Future work is planned to develop an ambulatory CEN recording device that could provide immediate notification of autonomic nervous system activity changes that might indicate autonomic dysregulation in healthy subjects and clinical disease states.

## Introduction

The autonomic nervous system (ANS) links the central nervous system (CNS; brain and spinal cord) with peripheral organ systems, including the integumentary (sweat glands), circulatory (heart, blood vessels), digestive (gastrointestinal tract glands and sphincters, kidney, liver, salivary glands), endocrine (adrenal glands), reproductive (uterus, genitals), respiratory (bronchiole smooth muscles), urinary (sphincters), and visual (pupil dilator and ciliary muscles) systems ^1-3^. The autonomic nervous system is colloquially divided into two main divisions: the “sympathetic” and “parasympathetic” nervous systems. However, both branches continuously coordinate through concerted feedback mechanisms to carefully control peripheral organ systems ^4,5^. A large body of empirical evidence suggests that autonomic nervous system imbalance is associated with various pathological conditions that can include heterogeneous disease states, such as diabetic autonomic neuropathy, hyperhidrosis, orthostatic intolerance/postural tachycardia syndrome, pure autonomic failure, autonomic dysreflexia, Takotsubo cardiomyopathy, and vasovagal syncope, and it also contributes to pathophysiology associated with autoimmune inflammatory disorders such as Rheumatoid Arthritis ^6,7^. Moreover, mental health disorders (e.g., Post-traumatic Stress Disorder (PTSD) and Major Depression Disorder) regularly exhibit circadian autonomic dysregulation, with heightened sympathetic and concomitant low parasympathetic drive most commonly reported ^8-12^.

The human cervical spine (neck) is the site of a confluence of autonomic neural structures that are in close proximity to each other, including the major parasympathetic neuronal output transmitted by the vagus nerve. The vagus nerve communicates directly to the visual, heart, respiratory, and digestive systems, and the major sympathetic neuronal output transmitted by the middle and superior cervical ganglion is located approximately 1-2 cm deep to the vagus nerve. The sympathetic ganglion, carotid body, and the vagus nerve are all localized within the carotid artery sheath, and, potentially due to this close proximity, sympathetic fibers have been observed in vagus nerve fascicles (called hitch-hikers), which further indicates multi-sourced neuronal signaling at this cervical level ^13^. The superior cervical ganglion and the thoracic sympathetic ganglion output directly to the integumentary, visual, circulatory, respiratory, and digestive organ systems. Given the immense peripheral organ system control generated from cervical autonomic neuronal structures found within the superficial cervical neck ^14^, decoding these signals to understand the role they play in health and disease could have significant impact on a host of conditions. Prior preclinical work has recorded resting microelectrode vagus nerve action potentials during lipopolysaccharide or inflammatory cytokine injection ^15-17^. Recent work has shown that vagus nerve action potentials uniformly synchronize with the respiratory cycle in porcine ^18,19^ and in one recent human microelectrode study ^20^. Other preclinical work measured (via cuff electrode) superior cervical ganglion activity with hypertensive stress tests, i.e., injection of adrenaline ^21,22^ or during painful stimuli ^23^. These studies uniformly demonstrate immediate (within seconds) change in cervical sympathetic neuronal (superior cervical ganglion) activity with each challenge ^21-23^. However, the invasive implantation and the risk associated with acute and chronic surgical cuff electrode complications have likely precluded, to date, any reported human cervical vagus nerve, carotid body, or superior cervical ganglion recording with validated stress models.

To enable recording with human autonomic stress models, a noninvasive, adhesive-integrated and skin conformal silver-silver chloride electrode array was developed that is capable of conformal positioning over the human left superior anterior cervical area overlying multiple neural structures (i.e., the vagus nerve and its branches, trigeminal nerve branches, sympathetic chain and its branches, the hypoglossal and glossopharyngeal nerves, as well as muscle and dermal sympathetic nerves), and its ability to monitor cervical nerve activity was tested using two widely used and validated stress tests, the cold pressor stress test (CPT) and a timed respiratory challenge.

CPT is performed by immersing the hand into a container filled with ice water, which is known to trigger a sympathetic reaction that involves blood vessel constriction and, thus, an increase in blood pressure ^24,25^. It also increases the reflexive modulatory vagal tone by activating multiple brain stem nuclei that coordinate afferent and efferent vagus nerve signaling ^26,27^. The effect of CPT – namely pain – on heart rate is bimodal: subjects routinely demonstrated either an increase or a decrease ^28,29^. Likewise, in timed respiratory challenge studies, muscle sympathetic nerve activity was bimodal; subjects either increased or decreased muscle sympathetic nerve firing ^30^. To date, there is a paucity of inter-challenge within subject physiological measures with CPT and timed respiratory challenge. To fill this gap, the newly developed flexible adhesive electrode array was deployed for non-invasive cervical electroneurography (CEN) during a sequential CPT and timed respiratory challenge. Multi-stress challenge CEN measures could help to further disambiguate human autonomic biotype amongst healthy and disease states in several ways: 1) by facilitating the development of biomarkers of response to pharmacologic and or neuromodulation therapies, 2) enabling the prediction of inflammatory response to pain, and 3) aiding differentiation of sterile vs. non-sterile inflammation.

## Results

The state-of-the-art custom surface electrode array that was tested is adhesive integrated and flexible, so that it can be non-invasively attached to a subject’ s anterior cervical neck. The electrode array was placed lateral to the trachea and medial to the sternocleidomastoid for this study. Silver-silver chloride was utilized as the material of choice for the custom electrodes (**Figure 1**) in tandem with a customized biopotential data acquisition unit (**Figure 2c**). The custom electrode array allowed subjects to freely move (lateral rotation as well as forward, reverse, and lateral flexion and extension) without distorting the adhesiveness and robustness of the physical structure of the electrode array (**Figure 2a, 2b**). Power line noise and its harmonics were minimized by the low impedance between the electrodes and skin.

**Figure 1.**
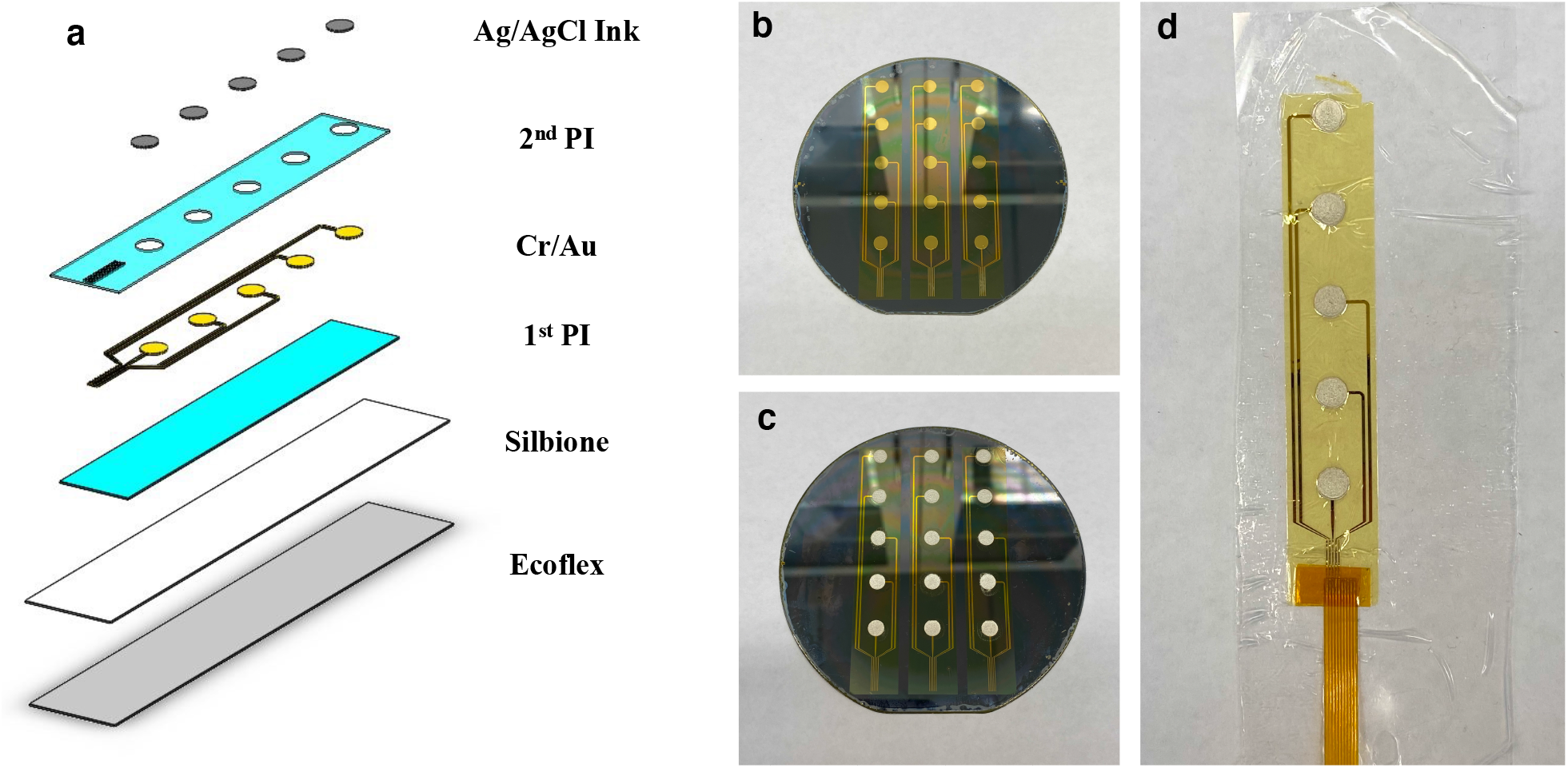
State-of-the-art custom surface electrode array for non-invasive testing of cervical neuronal activity. **(a)** Device cross section (5 mm sensor diameter and 500 um interconnect width). (**b)** Cleanroom post-processed wafer ready to be printed with Ag/AgCl ink. **(c)** Ag/AgCl ink screen-printed wafer on the active electrode region. **(d)** Device transfer-printed onto a self-adhering flexible silicone substrate (Ecoflex/Silbione) on PET backing via water soluble tape.

**Figure 2.**
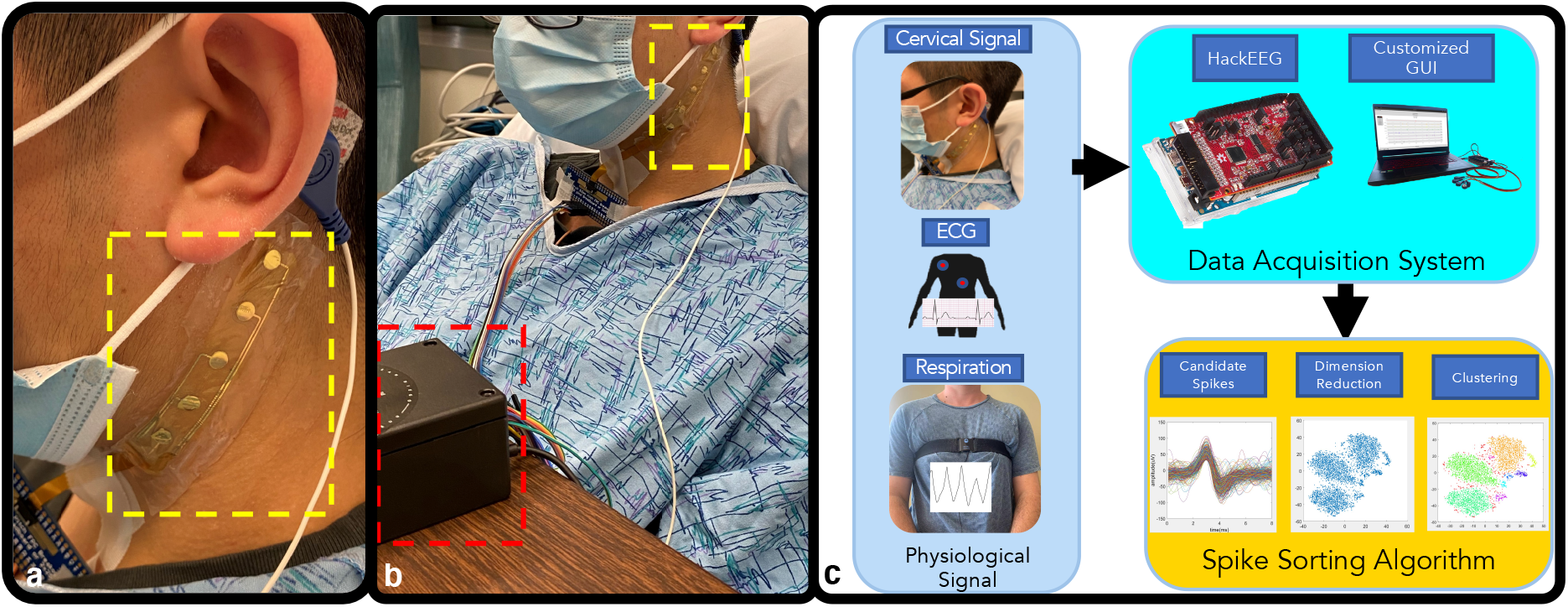
Custom design allows free movement without distorting the adhesiveness and robustness of the physical structure of the electrode array. **(a)** The custom surface electrode array is adhesive integrated, flexible, and non-invasively attached to the subject’ s anterior cervical neck, placed lateral to the trachea and medial to the sternocleidomastoid. A 3M RedDot electrode was used as a reference electrode. The four channels were aligned rostral to caudal, so that each channel was positioned over an anatomical target: 1) channel #1 over the nodose ganglion, 2) channel #2 over the nodose ganglion, 3) Channel #3 over the carotid artery (upper segment), and 4) channel #4 over the carotid artery (lower segment). **(b)** Data collection was carried out with flexible multi-electrodes attached to the wireless biopotential data acquisition board, i.e., the HackEEG (red dashed outline), and 3M RedDot deployed as ground/reference electrodes. **(c)** Subject recording pipeline and workflow. Cervical electroneurography, electrocardiography, and respiration were recorded; signals recorded to the data acquisition system underwent post processing algorithms.

Cervical electroneurography recordings were carried out with four electrode “channels” positioned rostral to caudal to evaluate cervical signal: 1) pre-to-post CPT and 2) during a timed respiratory challenge. All channels were run through a spike sorting algorithm to identify putative action potentials or nerve firing patterns associated with different nerve branches. Heart rate was derived from ECG recording by finding QRS peaks.

In a sample subject, the change in cervical neural firing was observed to coordinately increase with onset of CPT; the cervical neural firing was extrapolated from the spike sorting algorithm **(Figure 3a)**. Simultaneous recordings from Channel #1 (rostral overlying nodose ganglion, C2/3 cervical dermatome, and auriculotemporal branch of the trigeminal nerve) and heart rate were compiled **(Figure 3a)**. Amongst an array of clusters, responsive clusters were identified. Responsive clusters were defined as clustered groups that significantly increased in firing (by greater than 2 standard deviations) during CPT (for at least 40% of the stress challenge) compared to pre-CPT baseline activity.

**Figure 3.**
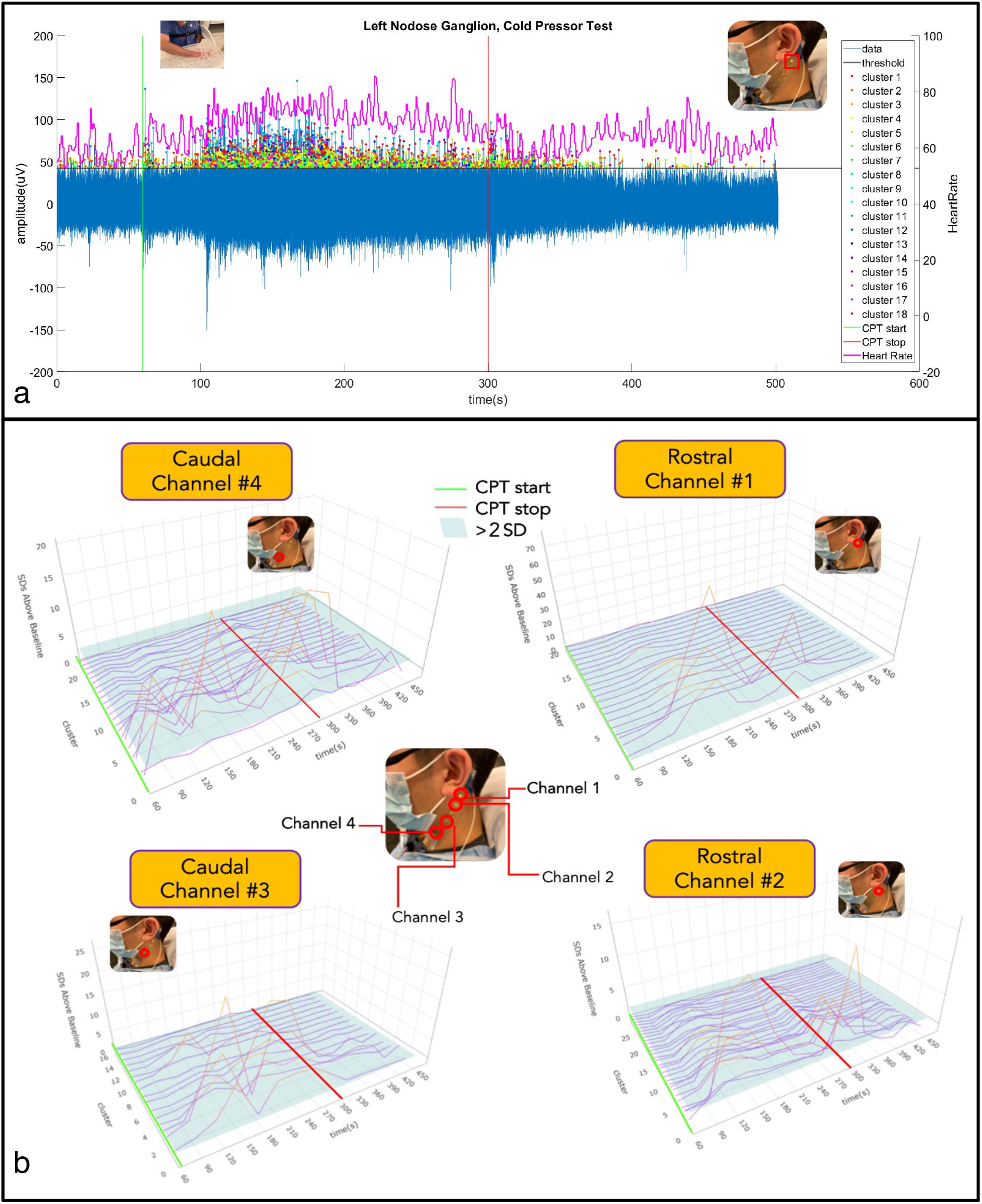
Cervical neural firing coordinately increases with onset of CPT. **(a)** Cervical Electroneurography (CEN) rostral Channel #1 recording post-spike sorting analysis for the cold-pressor test (CPT). The green vertical line is when the CPT started and the subject immersed their right hand into the ice-water bucket, while the red vertical line is when the CPT stopped. Channel #1 (blue vertical line = neural firing) captures the amplitude of the recorded neural potential data. The peak of each detected spike (blue line) that exceeds the threshold value was sorted by dots with different colors, representing the different clusters. Heart rate in beats per minute is plotted in magenta using the right y-axis. **(b)** Exemplar subject cervical electroneurography (CEN) neural firing change during the cold pressor test. <u>Top left</u>: Responsive clusters at the Caudal Channel #4 (located over the carotid artery, vagus nerve, glossopharyngeal nerve, sympathetic chain, and sensory C2/C3 dermatomal nerves) immediately increase in firing frequency with cold pressor test (CPT) onset (i.e., hand placed in ice water bath). <u>Top right</u>: Responsive clusters from the rostral channel #1 (located over nodose ganglion and in close approximation to the auriculotemporal nerve) demonstrate delayed firing onset (at 120–150 sec). <u>Bottom panels</u>: Both Caudal channel #3 (overlying the carotid artery, vagus nerve, glossopharyngeal nerve, sympathetic chain, and sensory C2/C3 dermatomal nerves) and Rostral channel #2 (located over nodose ganglion and in close approximation to the auriculotemporal nerve) increase in firing frequency at 90–120 sec post CPT initiation. In all panels, the transition from purple to orange lines indicates increases in cervical neural firing greater than 2 SD above baseline (i.e., prior to CPT start), and the green line denotes initiation of CPT, while cessation is denoted by the red line.

With the CPT challenge, approximately 20 unique spike clusters were recorded in each subject. Spike cluster firing activity was evenly divided into 5 temporally separated intervals over the duration of the CPT (0–20%, 20–40%, 40–60%, 60–80%, and 80–100%) with the aim of meticulously measuring onset and acquiescence of neural firing. Percent duration was employed to allow for temporal normalization of measures during the CPT. Average firing change (normalized by percentage change) for each channel across all clusters was computed at each CPT time interval and compared as follows: 1) within channel to baseline firing and 2) between channels at each time interval (**Figure 4**). All four channels at both the 0–20% and the 20–40% intervals showed significant increased firing activity when compared to baseline (p < 0.05). During the 0–20% CPT interval, Channel #4 (lower carotid artery) demonstrated greater neural firing (p<.05) when compared to both Channel #1 (over nodose ganglion) and Channel 2 (over lower nodose ganglion and 3rd trigeminal nerve, i.e., the auriculotemporal branch) as shown in Figure Furthermore, during the 20–100% intervals of CPT, the increase in overall firing was observed to be greater for Channel #2 than for Channel #1 (**Figure 4 b–d**).

**Figure 4.**
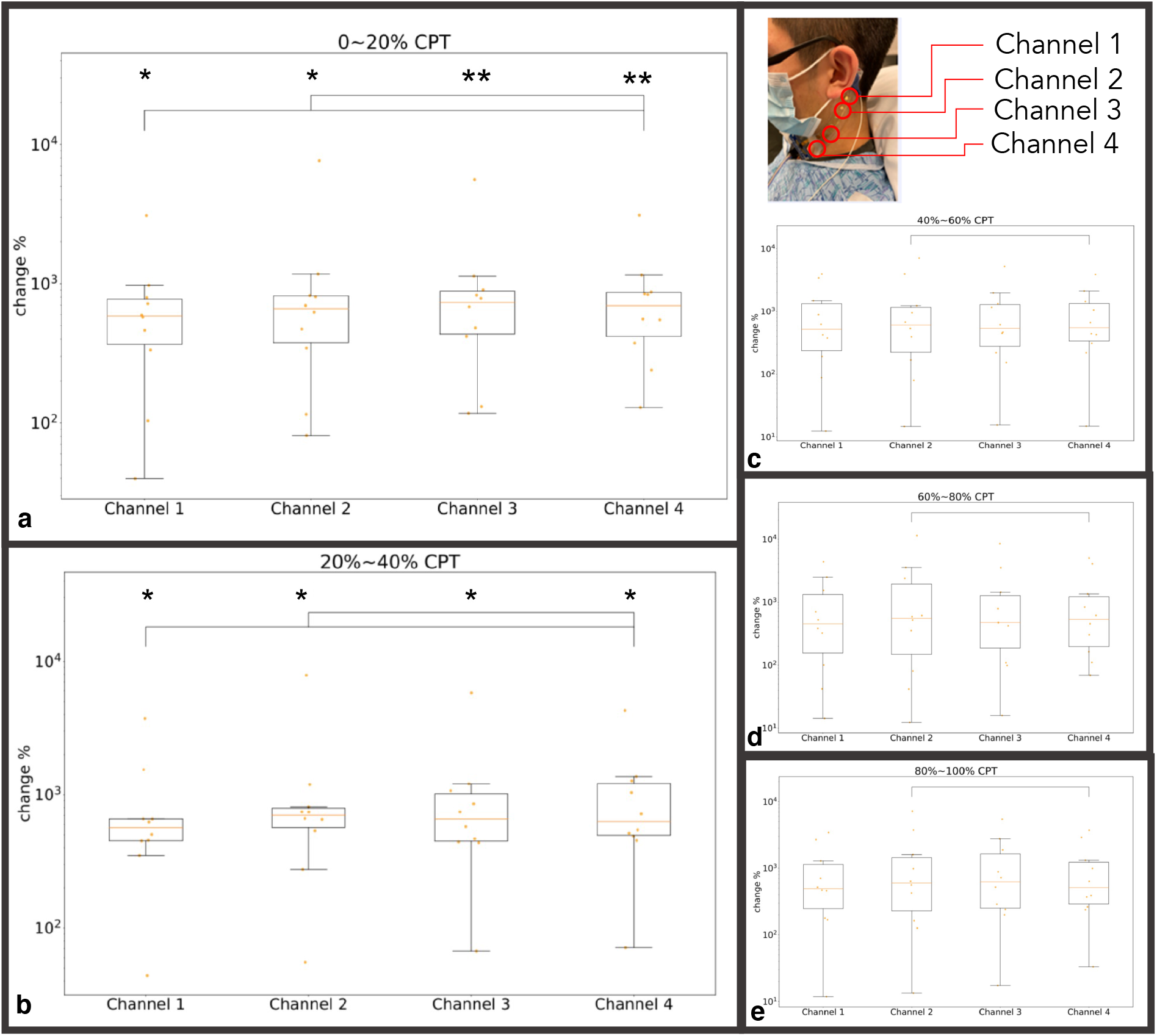
All subject channel activity across CPT indicated as % time intervals: **(a)** 0–20% of CPT; **(b)** 20–40% of CPT; **(c)** 40–60% of CPT; **(e)** 60–80% of CPT, and **(e)** 80–100% of CPT. Panel A: For the first interval of 0– 20%, channels #3 and #4 demonstrate highly significant increases in firing frequency (** = p<.001), while channels #1 and #2 also increased in firing frequency, but to a lesser extent (* = p<.05) when compared to baseline. Channel #4 (lower carotid artery) demonstrated significantly greater (brackets) firing than channels #1 and #2 (p<.05). Panel B: All channels remained at relatively increased activity (p<.05) when compared to baseline. An increase in firing in Channel #2 was observed when compared to channel #4 (p<.05). Panels C, D, E: All channels do not show increases in activity when compared to baseline. Channel 2 continued to demonstrate significantly greater firing than Channel 4 for the remaining CPT duration (p<.05). Channel #1 = overlying nodose ganglion; channel #2 = overlying lower nodose ganglion; Channel #3 = overlying carotid artery (upper segment); Channel #4 = overlying carotid artery (lower segment). X Axis: Channel 1–4; Y Axis: Percentage Increase in neural firing frequency with respect to baseline. activity.

Prior work by Mourot and colleagues ^29^ categorized subjects into two separate biotype groups, Cold Pressor Test Increaser (CPTi) or Cold Pressor Test Decreaser (CPTd), based on their HR responses under CPT (see methods). In this CPT experiment, similar group/biotypes to those described by Mourot and colleagues ^29^ were observed that were also equally distributed between CPTd (N=5) and CPTi (N=4). Building on Mourot’ s biotype categorization (i.e., groups separated based on CPTd or CPTi), we aimed to determine if nested biotypes could be observed during the CPT as well as during the timed respiratory challenge. During the CPT, greater neural firing (p<.05) was observed in the CPTi compared to the CPTd group for the first two intervals (0–20% and 20–40%) and across all channels (**Supplemental Figure 1**). The spline generalized estimating equations (GEE) showed that the change in firing frequency from 0 to 50 seconds was different between two CPT groups at Rostral Channel #2, Caudal Channel #3, and Caudal Channel #4 at the trend level (p<0.1) (**Supplemental Table 1, Supplemental Figure 2**). The models also show that the slope change in firing frequency from 0 – 50 seconds to 50 – 200 seconds was met a trend level in between CPT groups across all four channels (p<0.06) (**Supplemental Table 1**). Moreover, biotype-specific (CPTd vs. CPTi) changes in average CEN firing were observed over the duration of the timed respiratory challenge. Specifically, neural firing during pre-respiratory (initial 60 sec) challenge was compared to neural activity during post-respiratory challenge (post 60 sec) between groups (CPTd vs. CPTi). Significant increases in CEN firing was observed across all 4 channels during the timed respiratory challenge, while the inverse was observed in the CPTd group, which showed decreases in CEN firing over the timed respiratory challenge with significant changes noted in channel 4 **(Figure 5)**. Differences in CEN change pre-to-post timed respiratory challenge were identified between groups (CPTd vs. CPTi) across all channels, with the greatest change noted in Channel 1 **(Figure 5c)**. GEE analysis showed significant group effect on pre-to-post respiration neural firing change across all channels (p<.02; **Supplemental Table 2**). Moreover, in addition to the observed group effect on neural firing (both during CPT and timed respiratory challenge), there was a significant group effect (CPTd vs. CPTi) on heart rate during the timed respiratory challenge **(Supplemental Figure 1a)**.

**Figure 5.**
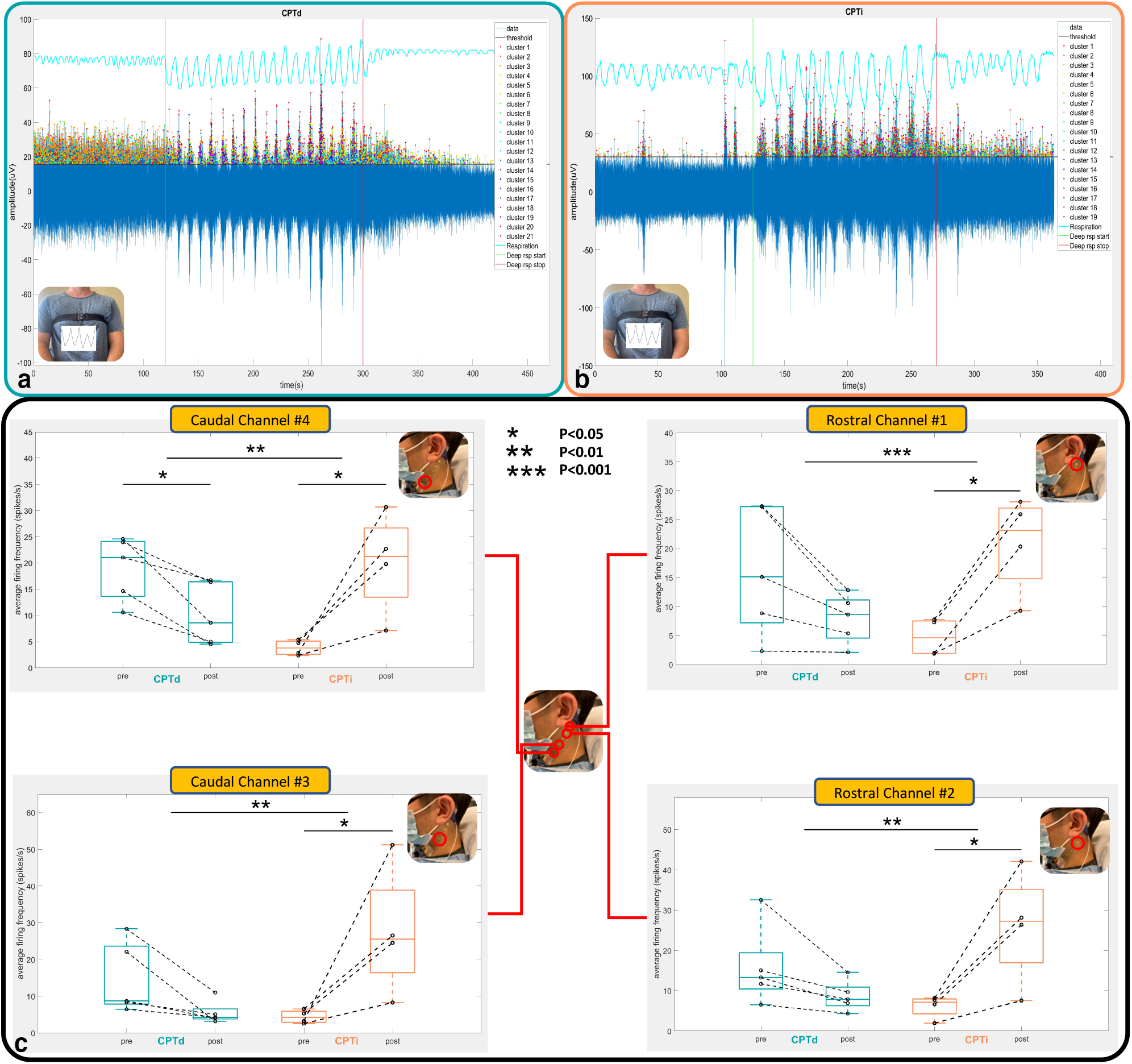
Neuronal firing comparison between CPTd and CPTi groups during the respiratory challenge. **(a)** and **(b)**: The green vertical line is when the respiratory challenge started, while the red vertical line is when the respiratory challenge stopped. The blue line captures the amplitude of the recorded electrode data for electrode positioned over the LNG (Channel #1). The peak of each detected spike (exceeding the threshold value) was also marked in dots with different colors in the positive y-axis, representing the different clusters to which they belong. The respiration belt voltage is plotted in cyan using the right y-axis. Exemplar CPTd CEN neural firing change during deep respiratory challenge. **(b)** Exemplar CPTi CEN neural firing change during deep respiratory challenge. **(c)** Average firing frequencies comparison between pre- and post-challenges for CPTd and CPTi groups across four channels. All channels in the CPTi group have significant neuronal activity increase after deep respiratory challenge (p<.05), whereas only Channel #4 in the CPTd group has significant decreased activity. Nevertheless, there are significant differences (p<.01) between the CPTd and CPTi groups on neuronal firing change across all channels.

## Discussion

In this study, a flexible, adhesive-integrated electrode array was presented for the non-invasive monitoring of neural activity at the anterior cervical area. The peel-and-stick device substituted the use of an invasive and cumbersome needle electrode. Because of the self-adhering and non-invasive design of the electrode array, the array enabled the measurement of electrical signals over a prolonged period. Overlying the electrode array at the left superior anterior cervical area spanning rostral to caudal in a diagonal fashion enabled measurement of biopotentials from the skin surface that could arise from the activation of multiple superficial and deep neural structures during CPT and a timed respiratory challenge. The nerve structures that could be measured by the electrode array include the vagus nerve and its branches, the sympathetic chain of interconnected ganglia, the hypoglossal nerve, and glossopharyngeal nerves, as well as muscle electric signals and the potential activity of dermal sympathetic nerves. These neural recordings provided a sufficient signal to noise ratio to enable an electrocardiographic-derived heart rate estimate and neuronal spike sorting throughout the CPT. Moreover, spike sorting of signals collected from the different channels of the electrode array revealed statistically significant temporal and spatial relationships. One basic requirement for spike sorting is the spatial relationship between channels. In the presented electrode device, both the electrode size and associated impedance, as well as the electrode shape and its arrangement on specific patterns can be customized ^31^. This versatile electrode shaping can aid in neural structure source identification. Accurate CEN potential sourcing of underlying nerval structures may provide a powerful tool to evaluate sympathetic versus parasympathetic activity and their correlated homeostatic processes. Once these signal sourcing challenges can be resolved, CEN may prove a valuable tool for assessing body homeostasis.

For the electrode design, Ag/AgCl was chosen as the sensor layer material due to high signal-to-noise ratio, lower skin-to-electrode impedance, and the non-polarizable nature of Ag/AgCl electrodes, which allows the Cl^-^ ion to take part in free charge exchange, preventing charge buildup _33-35_. Thin film technology enabled a high-resolution interconnect array that allows us to keep the form factor of the connector cable with 0.50 mm width (standard width used for a zero insertion force (ZIF) connector interface). The sensors were built using standard microfabrication processes followed by screen printing of Ag/AgCl ink, which yielded a final product that combined the best properties of each fabrication method (thin film and thick film). The thick-film technology enabled delivery of micrometer-thick Ag/AgCl ink that otherwise would not have been possible to deposit in a thin film process. Previously, it was demonstrated that flexible electrodes utilizing Ag/AgCl were successful in measuring other biopotential signals, such as electrogastrogram (EGG)^34^. Taken together, the developed Ag/AgCl electrode enabled a high-fidelity, low noise recording for this study’ s application.

The processing pipeline and electrode design were constructed to enable convenient and accurate neuronal recording. Specifically, an open-source HackEEG data acquisition unit was integrated with a Texas Instruments ADS1299 chip, which can sample 8 channels of data at 16k sampling rate with 24-bit ADC resolution. Because the neural activity targeted in this study is less than 1,000 Hz ^32^, an 8k Hz sampling rate was implemented, which efficiently enabled accurate reconstruction of critical spike neuronal waveforms. During measurements, the customized GUI was able to plot recordings in real time to monitor responses and abnormalities. Together, monitoring via the HackEEG biopotential board and custom GUI with our novel electrode array allowed for convenient recording of the subjects’ CEN. A future next step is to interface the flexible electrode array with the necessary remining hardware to amplify, digitize, and wirelessly broadcast the neural signals recorded, thus allowing ambulatory use.

The electrode design, recording technique, and electrode placement enabled the measurement of multiple neuronal signals characteristic of different possible sources. As in prior work, dermal sympathetic nerve firing, also referred to as skin sympathetic nerve activity, was ultimately detected, which is known to immediately increase with CPT onset; specifically, dermal sympathetic sudomotor firing results in eccrine sweat gland excretion that increases skin conductance, a phenomenon also known as electrodermal activity (EDA) ^32,36^. Other sources of signals likely were detected as well, including deeper muscular sympathetic neurons, carotid bodies, sympathetic chain, and ganglia spanning the carotid body, middle and superior cervical ganglion, in addition to action potentials emanating from the glossopharyngeal nerve, the vagus nerve and its rostral ganglia, the nodose and jugular ganglia.

Spike sorting the electrode array CPT recordings enabled identification of key temporal and spatial signal characteristics that could reflect different neural source responsivity to the CPT. As part of the spike sorting analysis, the detected spike waveforms were decomposed from each measurement into multiple clusters. After applying spike sorting to the measured signal, it was found that amongst all channels (Rostral to caudal Channel #1 = overlying nodose ganglion, Channel #2 = overlying lower nodose ganglion, Channel #3 = overlying carotid artery (upper segment), Channel #4 = overlying carotid artery (lower segment)), multiple neural clusters had significant increases in firing compared with baseline during CPT, followed by a gradual return to baseline after CPT termination (**Figure 3b**). Specifically, the neuronal firing was most significant (p < 0.05) during the first two (0–20% and 0–40% periods) of the five CPT time intervals **(Figure 4)**. Importantly, this detected temporal distribution of increased activity, demonstrating that the presented pipeline could detect (within 1 min) increases in cervical neuronal activity known to occur with CPT ^24-28^, which is indicative of changes in autonomic nervous system activity. Additionally, this activity pattern followed the expected effects of CPT on several neuronal structures within the coverage of the anterior CEN array, including the vagus nerve (increased vagal tone during CPT) ^25,26^, muscle and skin sympathetic nerve activity or EDA (increased activity followed by return to baseline), and carotid bodies (responsive to CPT induced increases in blood pressure)^27,32^. Collectively, the presented electrode array enabled the non-invasive detection and cluster isolation of neuronal spikes corresponding to CPT that may emanate from an aggregation or component of the vagus nerve, dermal, skin, muscle, and carotid body sympathetic response, while other neural structures may also contribute.

Unique to this study, significant spatiotemporally distinct cluster responsivity was observed when comparing Channels #1 and #2 (overlying the rostral auriculotemporal nerve and nodose ganglion) to Channels #3 and #4 (overlying the caudal carotid artery and sympathetic chain ganglia). Specifically, during the first 0–20% interval of the CPT, the caudal carotid artery sensor (Channel #4) demonstrated significantly higher firing frequency than the rostral nodose ganglion sensors (Channels #1 and #2) (p < .05) (**Figure 4**). In contrast, during the last 3 intervals of the CPT, the rostral nodose ganglion sensor (Channel #2) showed significantly higher firing frequency when compared to that of the caudal carotid artery sensor (Channel #4). Moreover, both caudal carotid artery sensors (Channels #3 and #4) demonstrated highly significant (p < .001) increases in firing frequency during the first interval of 0–20%, while the rostral sensors (Channels #1 and #2) also captured increased firing, but to a lesser degree than the caudal sensors (Channels #3 and #4) (p < .05) (**Figure 4**).

Prior work provided a premise for the observed delayed activity at the rostral Channels #1 and #2 (positioned over the trigeminal auriculotemporal branch distribution) when compared to the caudal Channels #3 and #4 (positioned over the carotid artery, carotid body, and sympathetic chain). In experimental studies of motion sickness ^37-39^ and virtual reality (VR) experiments that looked at “cybersickness” ^38^, immediate extremity dermal sympathetic neural activity was regularly observed; *however, increases in forehead (Trigeminal Nerve distribution) dermal sympathetic neural activity were delayed by several minutes or until coinciding nausea symptomatology began* ^37-39^. Vagus afferent and efferent signaling are well established as key mediators of nausea and vomiting ^40,41^, ^41,42^. Additionally, previous studies have shown coordinated decreases in vagus mediated high frequency heart rate variability (HRV) with the onset of nausea symptomatology in motion sickness, whereas the onset of strong nausea was proceeded by bursting increases in high frequency HRV ^43-45^. Likewise, vagus mediated high frequency HRV measures highly correlate with motion sickness/cybersickness onset with VR gameplay ^46^. High frequency HRV is often used as a proxy for vagus nerve activity and its effects on parasympathetic output ^47-49^ and is supported by cuff electrode vagus nerve firing correlates of HF HRV, while the reciprocal, i.e., inverse/negative correlation of vagus cuff firing to LF HRV, has not been reported to date ^50^. Furthermore, electrogastrography (EGG) measures slow wave myoelectric activity in the stomach that is directly dependent on vagus nerve signaling ^51^ and decreases with motion sickness/cybersickness onset ^52,53^. Our group showed that gastric antrum stimulation decreases EGG slow wave power ^54^, which is in line with recent pre-clinical work by Tan et. al ^55^. Moreover, our group showed brainstem, lower medulla, dorsal motor nucleus of the vagus (DMX), nucleus tractus solitarius (NTS), and rostral ventrolateral medulla (RVLM) blood-oxygen-level dependent (BOLD) activation with thermal pain ^56^. Recently, others have now confirmed these specific brainstem activation patterns in numerous experimental pain paradigms, such as mechanical ^27^, thermal ^56^, and CPT ^57^. Remarkably, these brainstem areas that coordinate complex autonomic neural circuitry, including vagal parasympathetic (DMX, NTS) and sympathetic nuclei (RVLM), demonstrated a relative delay in onset activation (similar to delays observed with motion and cybersickness) when compared to cortical clusters, including the supplementary motor and insula cortices ^57^. Collectively, these studies indicate that delayed changes in vagus nerve signaling, as well as specific dermatomal sympathetic neural activity “reflex circuits,” coordinate in tandem with onset of nausea in motion/cybersickness and during experimental pain paradigms such as CPT. In this study, these constructs are supported by the observation that caudal Channels #3 and #4 (positioned over the carotid artery, carotid body, and sympathetic chain) immediately (0–20% CPT interval) increased in neural sympathetic activation, while Channels #1 and #2 (positioned over the trigeminal auriculotemporal branch distribution) showed relatively less activity at this interval. However, greater activity in Channel #2 was observed in the last half of the CPT. As sympathetic output to the forehead is controlled by the Trigeminal nerve, the relative delay in neural firing in Channels #1 and #2 (positioned over the auriculotemporal branch of the Trigeminal nerve and the nodose ganglion) coincides with previous reports that demonstrate a delayed increase in trigeminal nerve sympathetic output that is known to occur with vagal-mediated reflex stress responses, associated with onset of motion/cybersickness, and CPT ^37-39^. Future work will measure EGG, magnetoencephalography (MEG), and cervical electroneurography (CEN) during CPT to record multi-sourced coordinate neural changes that may further identify high resolution temporally dependent autonomic reflexes and give insight into visceral-brain coupling recently observed between EGG and MEG recordings ^58^.

Distinctive to this study, subjects underwent two sequential autonomic stress tests, i.e., the timed respiratory challenge followed by the cold pressor test, while simultaneously recording heart rate and CEN change in neural firing. Human defense responses covary with autonomic regulatory response subtypes; individuals regularly demonstrate distinct autonomic biotypes with noxious stress challenge tests such as cold pressor (i.e., CPTd and CPTi) ^25,59-65^. In this study’ s cohort, an approximately equal number of subjects were categorized as CPTd (N=5) and CPTi (N=4). Extraordinarily, biotype-specific change in average CEN firing during the CPT and timed respiratory challenges was consistently observed (**Figure 5, Supplement Figure 1, 2**). To our knowledge, this is the first cross challenge report that indicates a direct relation between timed respiratory challenge CEN and heart rate change during CPT. Prior work by our group demonstrated that pre-deployed warfighter dysautonomia is predictive of eventual development of post-traumatic stress disorder (PTSD) in servicemembers ^66^. Because PTSD is known to have a baseline hyper-sympathetic drive, it is concordantly co-morbid with cardiac disease risk ^67^. Therefore, it was postulated that the hyper-sympathetic timed respiratory challenge response identified in the CPTi group may indicate a biotype at risk for mental health disorder and or hyperinflammatory response. In support of this construct, PTSD sympathetic neural firing is highly increased during CPT when compared to healthy controls ^68^. Future work will determine if CEN-derived hyper-sympathetic biotypes can predict immune cell response in an in-vivo human inflammatory model (intravenous lipopolysaccharide injection), in healthy subjects and servicemembers with PTSD.

Although these findings are promising, the present study has several limitations. First, the data are cross-sectional with relatively small sample size. This pilot study was meant to demonstrate feasibility of the flexible, adhesive-integrated electrode array for CEN recording during validated stress challenges, i.e., CPT and timed respiratory challenge. These findings need to be validated in a larger study cohort of healthy subjects and patients with a mental health disorder, namely PTSD. In addition, these findings would have more impact if the study were done in larger populations of both sexes, with the aim to identify sex effects. For instance, we did not control for menstrual cycle for all female participants, which could impact CEN recording during CPT ^68^. Prior work also demonstrated that sympathetic responses are both longer in duration and larger in amplitude during afternoon testing ^69^. Although we did not control for exact time of day for the timed respiratory challenge or CPT, all testing occurred within the window of 13:00 to 18:00 and not during nighttime hours.

In conclusion, this study revealed three major findings: **First**, a flexible, adhesive-integrated electrode array was shown to be capable of non-invasive monitoring of cervical neuronal activity (spike cluster firing) during two validated stress challenges, CPT and timed respiratory challenge. **Second**, significant spatiotemporally distinct sensor specific cluster responsivity was observed; rostral channels (overly trigeminal and vagus nerve branches) demonstrated less activity during the first portions of the CPT with a rebound increase during the last half of the CPT, emulating d neural activity observed in motion and cybersickness. **Third**, a potential biotype marker based on heart rate was identified across both the CPT and timed respiratory challenges. During both challenges, an increase in neural firing and heart rate was observed in the CPTi group, while the inverse was consistently observed in the CPTd group. In aggregate, the presented novel electrode array shows promise in this pilot trial for the non-invasive detection of autonomic nervous system change in activity (via CEN) across multiple validated stress challenges. More work is required to confirm CEN robustness and capability to derive biotypes in healthy control and patient populations.

## Methods

### Electrodes Array Fabrication

A 4” silicon wafer was cleaned with acetone, isopropyl alcohol (IPA), de-ionized (DI) water, and IPA again followed by drying with an N_2_ gun. The wafer then underwent a dehydration bake at 180°C on a contact hotplate. Subsequently, polydimethylsiloxane (PDMS), which acts as a weakly adhering substrate, was spun coated at 4000 rpm. Polyimide (HD Microsystems, Inc. - Parlin, NJ) was then spun coated at 4000 rpm followed by a soft bake at 110°C for 1 min and at 150°C for 5 min on a contact hotplate. A full hard bake in an N_2_-rich environment oven at 300°C then followed. Metalization was performed on an electron beam evaporator (Temescal BJD 1800 E-Beam Evaporator – Livermore, CA) to yield a 10 nm chrome and a 500 nm gold layer. Sensors (5 mm diameter) and interconnects (500 um width) were then defined via standard lithographic procedures. Another layer of polyimide was then applied using the same parameters as above. Photolithography and reactive ion etching (RIE) with O_2_ plasma followed to define the bottom and top polyimide layers, which acted as insulator layers.

Post cleanroom processing, a silver/silver chloride ink (Creative Materials, Inc. – Ayer, MA) was screen printed onto the active sensor areas via a standard stainless-steel stencil (Metal Etch Services, Inc. – San Marcos, CA). A custom zero insertion force (ZIF) connector was then interfaced with the electrode array using anisotropic tape (3M, Inc. – Saint Paul, MN) to facilitate bonding by applying heat and pressure on the bonding sites. Water soluble tape (3M, Inc. – Saint Paul, MN) was then used to transfer print the device from the silicon wafer to a flexible silicone substrate made on a 5” petri dish that was spun coated at 3000 rpm with Ecoflex (Smooth On, Inc. – Macungie, PA), which acted as a backbone silicone, and Silbione RT Gel (Elkem, Inc. – Brunswick, NJ), which acted as an adhesive silicone. The silicone substrate and the device were then peeled off and transferred to a thin sheet of polyethelene terephthalate (PET) (Grafix – Maple Heights, OH), which acted as a backing that eased handling of the device. The silicone bilayer was then cut into a rectangular shape, which yielded the final sensor array as shown in **Figure 1**.

### Participants

A total of 10 mentally and physically healthy subjects (2 females and 8 males, ages 21.8 ± 2.1) were recruited and enrolled in the study. The study was approved by the University of California San Diego Institutional Review Board (IRB#171154) and all research was performed in accordance with guidelines and regulations. All subjects provided written informed consent prior to participation and agreed to disclose identifying images for open-access publication.

### Electrode Placement

Conductive gel was applied to the electrode surface using a syringe to improve conductivity. The device was then applied to the neck with the rostral electrode approximately at the skin above the Trigeminal nerve auriculotemporal branch and the nodose ganglion and the caudal electrode at the skin above the carotid artery. Once attached, the PET backing was peeled off **(Figure 1a)**. The ZIF connector was attached to a breakout board (Adafruit Industries, Inc. NY, USA), which was then connected via cables to a biopotential data acquisition HackEEG board (Starcat LLC Seattle, WA) along with ground and reference electrodes (3M Red Dot, 3M, Inc – Saint Paul, MN). Electrocardiography (ECG) was also monitored by the 3M electrodes that were placed on the right upper chest and left bottom rib.

### Data Collection

Subjects were recruited at the University of California San Diego. Before the device application, the electrode placement area was abraded with abrasive pads (BIOPAC System Inc, Goleta, CA, USA) and prepped using alcohol pads to exfoliate the skin. A ground electrode was placed on the left forearm of the subject, and an additional reference cup electrode was placed ipsilaterally above the flexible sensor array at the mastoid. Data was collected and streamed to the open source HackEEG data acquisition system (Starcat LLC, Seattle WA, USA). The HackEEG is a high-performance, open-source Arduino Due shield for the Texas Instrument ADS1299 system on a chip. It can digitize 8-channel bio signals simultaneously at 24-bit analog-digital conversion resolution at 8,000 samples per second with user-selectable gain from 1 to 24. The HackEEG system is enclosed in a 3D printed box with the dimensions of 13 cm × 7 cm × 5 cm **(Figure 1b)**.

### Timed Respiratory Challenge

Respiration vital sign change was measured by a respiration belt transducer (BIOPAC system Inc, Goleta, CA, USA). The respiration belt was modified to function as a potentiometer, soldered in series with a 5k ohm resistor that was powered by the 3.3V power pin on the Arduino board. The voltage variation caused by the resistance change during each breath was measured by the HackEEG system as one of the biopotential channels. The respiration signal was filtered by a 5 Hz zero-phase low-pass filter in post-processing.

120 seconds after baseline recording, the subjects underwent a timed respiratory challenge. The timed respiratory challenge consisted of regular breathing (5 s inhale and 5 s exhale) that was carried out for 10 cycles. Post timed respiratory challenge data was then recorded for an additional 120 seconds. One subject’ s respiratory data was excluded due to irrevocable data loss.

### Cold Pressor Test

Five minutes after the timed respiratory challenge, subjects were asked to perform the CPT task. Subjects were required to fully immerse their right hand into an ice water container to determine the cold pressor tolerance. Chilled water and freezer fresh ice (3°C) were immediately added to the container with water temperature kept at 4–7°C. Cold pressor tolerance was carried out for each subject; subjects were asked to keep their right hand in the ice water container for at least 1 minute (5 minutes maximum) and only withdraw their hand if they could no longer tolerate the pain. Subjects were told to stay still as well as avoid talking and swallowing during the experiment to avoid EMG activity/artifacts.

### Defined Cold Pressor Test decreaser (CPTd) and Cold Pressor Test increaser (CPTi)

In this work, two common groups were identified: Cold Pressor Test increasers (CPTi) and Cold Pressor Test decreasers (CPTd). As previously defined by Mourot and colleagues ^29^, individuals in the CPTi group experienced increased HR when reacting to CPT, i.e., first an increase in HR was observed and then either a further increase in HR occurred, or the HR remained elevated until the end of the test. It should be noted that if HR decreased less than 5 beats, the HR was considered maintained, and the participant was categorized to the CPTi group. As previously defined by Mourot and colleagues ^29^, CPTd individuals were observed to react to the test with an initial increase followed by a decrease in HR of more than 5 beats per min (mean over 10 s) compared to the peak HR achieved during the test.

### Data Processing

A customized graphic user interface (GUI) installed on a Linux terminal was written in Python to monitor and display physiological signals in real time. In post processing, the cervical signal was filtered by a 20–1,000 Hz zero-phase bandpass filter. Powerline noise and harmonics were also removed by notch filters. In each cervical neural recording, a spike sorting algorithm was carried out to differentiate detected spikes into different clusters. First, spike detection was performed by setting a threshold as:

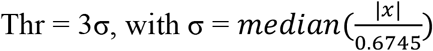

where x is the bandpass filtered signal ^70^. For each detected spike, a waveform of 80 samples (10 ms) was saved as the candidate waveform template, then aligned to their maximum at data point 30 (3.75 ms). Dimensionality reduction for feature extraction was run through T-distributed stochastic neighbor embedding (t-SNE) on all candidate spikes to enable a representative visualization ^71^. The low dimensional projection was then run through density-based spatial clustering of application with noise (DBSCAN), an unsupervised classification algorithm, ^72^ to cluster spikes into groups. The clustered neural groups were characterized by different firing rate and amplitude behaviors. The firing rate was counted as the number of neural group activity per second, and the amplitude was derived by calculating the peak-to-peak difference of each detected waveform **(Figure 1c)**.

### Statistical Analysis

Two-tailed paired sample t-test was used to identify differences between the firing frequency change of each individual channel when compared to baseline (Pre-CPT) activity across 5 intervals for the entire CPT duration (**Figure 4**), as well as between each channel at each time interval (**Figure 4**). The same methodology was used to test pairwise pre-to-post difference in 1) channel firing during the timed respiratory challenge (**Figure 5c**), 2) channel firing over the duration of the CPT challenge (**Supplementary Figure 1b**), and 3) HR change during the CPT (**Supplementary Figure 1a**), for all within and between CPTd and CPTi group comparisons.

Two tailed two-sample t-test was used to compare neuronal firing differences between the CPTd (N=5) and CPTi (N=4) groups during the timed respiratory challenge, i.e., average firing frequency difference between the first and last 60 seconds of measurement (**Figure 5c**). All post-processing and statistical analysis were carried out with MATLAB software (MathWorks Inc, MA, USA).

To further understand interactions between groups (CPTi vs. CPTd) neural firing over time on each channel, spline generalized estimating equations (GEE) ^73^ across the whole time series and with knots at the 50 sec and 200 sec time points were used. The spline GEE models regressed firing frequency on time, group, and time by group interaction (**Supplemental Figure Supplemental Table 1**).

To further understand interactions between groups (CPTi vs. CPTd) neural firing during the timed respiratory challenge, a linear regression model with inference based on GEE was utilized. Differences in groups (CPTi vs. CPTd) neural firing pre-to-post the timed respiratory challenge were determined with the generalized estimating equations (**Supplemental Table 2**).

## Acknowledgements

The authors would also like to acknowledge Marcelo Aguilar and Jamie Burks for their useful discussion.

## Author contributions

IL, TC proposed the experimental concept. IL, TC, YB and JK designed methodology and experiments. RR suggested experimental improvements. JK, AN, NS, TP, VW, BT, AS designed the surface electrode. BH designed graphic user interface. YB, JK, AN collected data. YB analyzed data. TW, XT suggested statical plan and conducted statistical analysis. IL, YB and JP wrote the primary content of the paper. All other authors participated in the editing of the final manuscript.

## Data availability Statement

Data is available at the authors’ discretion upon direct request to the corresponding author.

