## Supplemental Material for "Measurement of Cervical Neuronal Activity during Stress Challenge Using Novel Flexible Adhesive Surface Electrodes"

### Supplementary

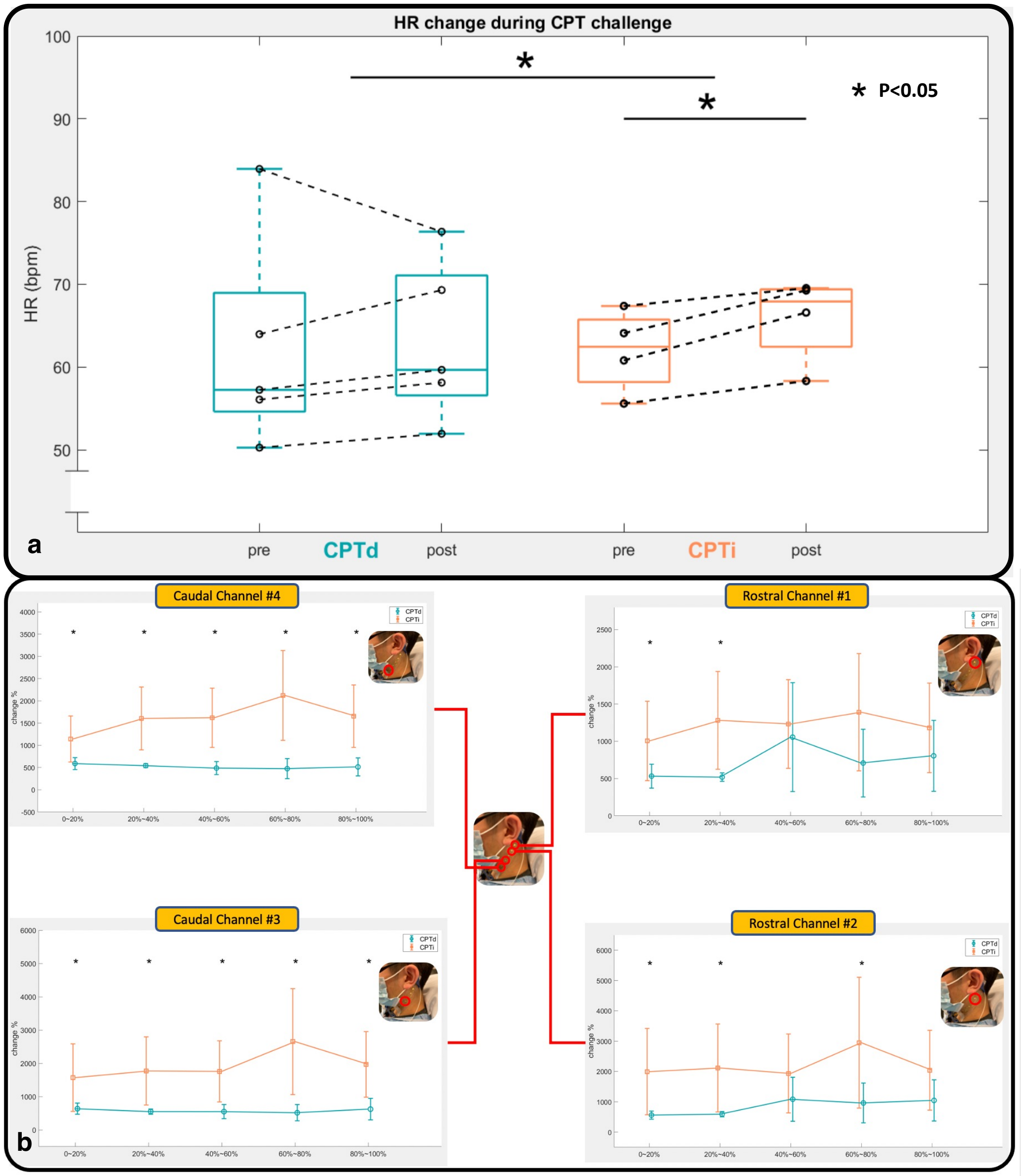


**Supplemental Figure 1. Heart rate comparison between pre and post timed respiratory challenge for CPTd and CPTi groups. (a)** The CPTi group had a significant change in HR after timed respiratory challenge (p<.05), whereas CPTd did not show a significant difference. Between group (CPTd vs CPTi) we observed a significant difference (p<.05) in HR change pre- to post timed respiratory challenge (p<.05). **(b) The comparison of average firing rate change with respect to baseline activity between CPTd and CPTi group across CPT intervals.** (Top Right) Rostral Channel #1 overlying the nodose ganglion and in close approximation to the auriculotemporal nerve only demonstrated a significant difference between two groups at the first two intervals, i.e., 0~40 % interval. (Bottom Right) Rostral Channel #2 overlying the nodose ganglion and in close approximation to the auriculotemporal nerve showed significant difference between two groups at 0~40% interval and 60~80% interval. (Left Panels): Caudal Channel #3 and Caudal Channel #4, overlying the carotid artery, vagus nerve, sympathetic chain and sensory C2/C3 dermatomal nerves, showed significant difference between CPTd and CPTi groups across the whole CPT challenge. *=p<.05

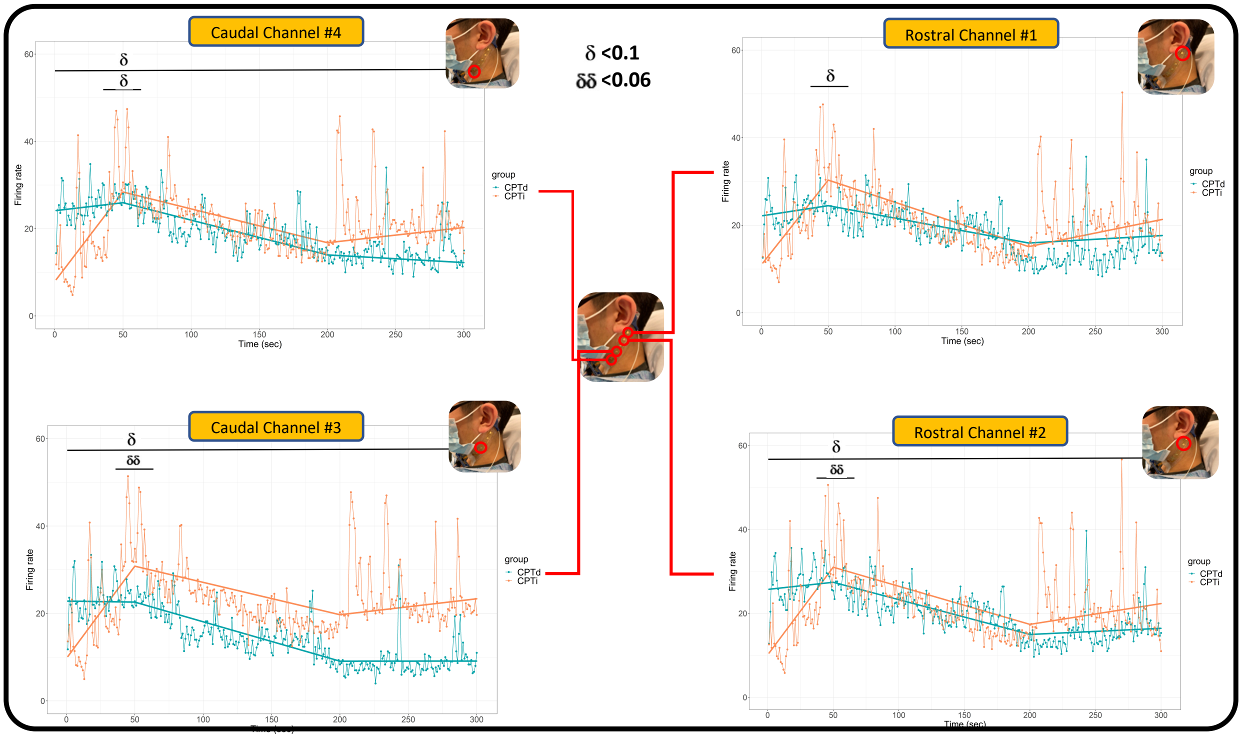


**Supplemental Figure 2. The average firing rate across the 5-minute CPT experiment between CPTd and CPTi groups along with the fitted line regressed by the generalized estimating equations (GEE). (Right panels)** Rostral Channel #1 and Rostral Channel #2 overlying nodose ganglion and in close approximation to the auriculotemporal nerve **(Left Panels)**: Caudal Channel #3 and Caudal Channel #4, overlying the carotid artery, vagus nerve, sympathetic chain and sensory C2/C3 dermatomal nerves. **δ**: significant trend level (p<0.1), **δδ**: significant trend level (p<0.06).

| | | Estimate | Std. Error | z-value | Pr(>\|z\|) |
|---|---|---|---|---|---|
| Channel 1 | (Intercept) | 22.13727 | 8.030872 | 1.736494 | 0.082477 |
| | Group | -11.1597 | 9.433433 | 1.108036 | 0.267846 |
| | Time | 0.046648 | 0.142015 | 0.325375 | 0.744897 |
| | sec50plus | -0.1035 | 0.129019 | 0.757586 | 0.448699 |
| | sec200plus | 0.073981 | 0.057867 | 0.923407 | 0.355795 |
| | Group:Time | 0.340105 | 0.197418 | 1.517496 | 0.129141 |
| | Group:sec50plus | -0.38456 | 0.172567 | 1.83062 | **0.067157** |
| | Group:sec200plus | 0.089454 | 0.081863 | 1.088715 | 0.27628 |
| Channel 2 | (Intercept) | 25.70524 | 8.216182 | 1.819041 | 0.068905 |
| | Group | -15.6124 | 9.212211 | 1.493718 | 0.135249 |
| | Time | 0.03404 | 0.123743 | 0.273111 | 0.784768 |
| | sec50plus | -0.11694 | 0.111537 | 0.944843 | 0.344739 |
| | sec200plus | 0.097491 | 0.071408 | 0.957679 | 0.338225 |
| | Group:Time | 0.384074 | 0.18975 | 1.711837 | **0.086927** |
| | Group:sec50plus | -0.39199 | 0.164356 | 1.923029 | **0.054476** |
| | Group:sec200plus | 0.042894 | 0.099004 | 0.441997 | 0.658492 |
| Channel 3 | (Intercept) | 22.82676 | 6.726084 | 1.867105 | 0.061887 |
| | Group | -13.1198 | 8.174923 | 1.431117 | 0.152397 |
| | Time | -0.00455 | 0.090682 | 0.050177 | 0.959982 |
| | sec50plus | -0.08555 | 0.092522 | 0.847293 | 0.396832 |
| | sec200plus | 0.090392 | 0.058062 | 1.029675 | 0.303163 |
| | Group:Time | 0.426751 | 0.195826 | 1.803727 | **0.071274** |
| | Group:sec50plus | -0.41072 | 0.174608 | 1.90857 | **0.056318** |
| | Group:sec200plus | 0.020131 | 0.082642 | 0.246157 | 0.805561 |
| Channel 4 | (Intercept) | 24.11242 | 7.419533 | 1.842093 | 0.065462 |
| | Group | -16.2104 | 8.835778 | 1.586886 | 0.112538 |
| | Time | 0.036527 | 0.095804 | 0.375909 | 0.706985 |
| | sec50plus | -0.11626 | 0.102424 | 1.005312 | 0.314746 |
| | sec200plus | 0.0622 | 0.062057 | 0.773313 | 0.439337 |
| | Group:Time | 0.373464 | 0.193448 | 1.651718 | **0.098592** |
| | Group:sec50plus | -0.37127 | 0.174747 | 1.775078 | **0.075885** |
| | Group:sec200plus | 0.050287 | 0.080741 | 0.636409 | 0.52451 |

**Supplemental Table 1. The fixed and the interaction effects of group and time on firing frequency from the generalized estimating equation.** Group: CPTd vs. CPTi. Time: Time in second as a variable over the entire 5-minutes session. Sec50plus: Inflection time point at 50 second. Sec200plus: Inflection time point at 200 s. There were significant CPT-group-by-time interaction effects from 0 to 50 seconds at Rostral Channel #2, Caudal Channel #3 and Caudal Channel #4, and significant CPT-group-by-50-second-inflection-time interaction effects across all four channels ($p<0.1$).

| | | Estimate | Std. Error | z-value | Pr(>\|z\|) |
|---|---|---|---|---|---|
| Channel 1 | (Intercept) | -8.26536 | 2.8317 | 1.775066 | 0.075887 |
| | Group×Neural Firing | 24.46228 | 3.822498 | 2.662324 | **0.00776** |
| Channel 2 | (Intercept) | -7.15148 | 2.50521 | 1.760316 | 0.078354 |
| | Group×Neural Firing | 27.07982 | 5.597999 | 2.504695 | **0.012256** |
| Channel 3 | (Intercept) | -9.27842 | 3.2602 | 1.758276 | 0.078701 |
| | Group×Neural Firing | 32.50936 | 8.351357 | 2.3398 | **0.019294** |
| Channel 4 | (Intercept) | -8.72743 | 1.847255 | 2.021129 | 0.043266 |
| | Group×Neural Firing | 24.96773 | 4.523209 | 2.586421 | **0.009698** |

**Supplemental Table 2. Summary result of the group (CPTd vs. CPTi) effect on pre-to-post respiration firing frequency change from the generalized estimating equation.** All four channels showed significant group effect difference ($p<0.02$).
